# Adaptive rescheduling of error monitoring in multitasking

**DOI:** 10.1101/2020.04.15.043059

**Authors:** Robert Steinhauser, Marco Steinhauser

## Abstract

The concurrent execution of temporally overlapping tasks leads to considerable interference between the subtasks. This also impairs control processes associated with the detection of performance errors. In the present study, we investigated how the human brain adapts to this interference between task representations in such multitasking scenarios. In Experiment 1, participants worked on a dual-tasking paradigm with partially overlapping execution of two tasks (T1 and T2), while we recorded error-related scalp potentials. The error positivity (Pe), a correlate of higher-level error evaluation, was reduced after T1 errors but occurred after a correct T2-response instead. MVPA-based and regression-based single-trial analysis revealed that the immediate Pe and deferred Pe are negatively correlated, suggesting a trial-wise trade-off between immediate and postponed error processing. Experiment 2 confirmed this finding and additionally showed that this result is not due to credit-assignment errors in which a T1 error is falsely attributed to T2. For the first time reporting a Pe that is temporally detached from its eliciting error event by a considerable amount of time, this study illustrates how reliable error detection in dual-tasking is maintained by a mechanism that adaptively schedules error processing, thus demonstrating a remarkable flexibility of the human brain when adapting to multitasking situations.

**Significance Statement:** Multitasking situations are associated with impaired performance, as the brain needs to allocate resources to more than one task at a time. This also makes it more difficult to detect one’s own performance errors in such complex scenarios. In two experiments, we recorded error-related electroencephalographic (EEG) activity and found that the commonly assumed fixed temporal succession of control processes in error monitoring can be strategically interrupted. Individual processes of error detection can be temporally rescheduled to after completion of competing tasks. This reduces interference between the neural task representations and supports a more efficient execution of concurrent tasks in multitasking.

## Introduction

Adaptive human behavior requires an error monitoring system that constantly checks whether an error has occurred and initiates adjustments to cognition and behavior accordingly. Numerous studies could show that such a system exists in the human brain, detecting errors fast and reliably (Ullsperger, Fischer, Nigbur, & Endrass, 2014). Most studies, however, investigated error monitoring when single tasks were executed in isolation. In contrast, everyday behavior is characterized by the concurrent execution of temporally overlapping tasks. As dual-tasking is associated with decrements in task performance (Pashler, 1994; Tombu & Jolicœur, 2003), it is plausible to assume that also error monitoring suffers under these conditions. Indeed, recent evidence suggests that dual-tasking leads to specific impairments to the neural correlates of error monitoring (Klawohn, Endrass, Preuss, Riesel, & Kathmann, 2016; Weißbecker-Klaus, Ullsperger, Freude, & Schapkin, 2016). In the present study, we investigate how the brain adapts to these dual-tasking conditions. By considering error-related brain activity in event-related potentials (ERPs), we ask how monitoring processes are reorganized to maintain reliable error detection when the brain is confronted with two temporally overlapping tasks.

Close temporal succession of more than one task leads to considerable interference between the subtasks, resulting in performance decrements compared to the execution of single tasks (Jersild, 1927; Telford, 1931). As dual-task interference arises predominantly in central, decision-related processes, the brain adapts to this interference by serializing these processes, granting attentional resources to only one task representation at a time (Logan & Gordon, 2001; Meyer & Kieras, 1997; Pashler, 1994). Neuroscientific studies suggested that this interference as well as the resulting serialization is linked to a fronto-parietal network that is able to grant access to focused attention and conscious control to only one task representation, by synchronizing distinct brain regions in a self-amplifying process. A considerable number of ERP studies links this serialization process to the P300 component (Dehaene, Kerszberg, & Changeux, 1998; Del Cul, Baillet, & Dehaene, 2007; Dell’Acqua, Jolicoeur, Vespignani, & Toffanin, 2005; Gross et al., 2004; Hesselmann, Flandin, & Dehaene, 2011; Sergent, Baillet, & Dehaene, 2005; Sergent & Dehaene, 2004; Sigman & Dehaene, 2008). For example, Sigman and Dehaene (2008) found in a dual-tasking paradigm that neural correlates of early attentional processes were executed in parallel for both tasks, whereas the P300 for the second task (T2) was delayed until response selection for the first task (T1), represented by its own P300, had been completed (see also Dell’Acqua et al., 2005; Hesselmann et al., 2011). Recently, Marti, King, & Dehaene (2015) provided evidence that this serialization of task selection is the result of a competition between task-related processes when they strive for access to a higher-level attentional workspace. Neural representations of the two tasks repel each other in a way that initially, only processing of T1 is enabled, and that of T2 is actively delayed.

It has recently been suggested that also processes involved in the detection and evaluation of performance errors suffer from dual-tasking interference. Research has robustly identified two ERPs associated with distinct stages of error monitoring, which are commonly reported to be executed in a fixed temporal cascade immediately following the erroneous response (Ullsperger, Fischer, Nigbur, & Endrass, 2014) – the error-related negativity (Ne/ERN, Falkenstein et al., 1991; Gehring et al., 1993) and the error positivity (Pe, Overbeek et al., 2005). The Ne/ERN, an early fronto-central negativity on error trials, has been assumed to reflect a mismatch between the correct and the actual response (Coles, Scheffers, & Holroyd, 2001), response conflict (Yeung, Botvinick, & Cohen, 2004), or prediction errors that accompany performance errors (Holroyd & Coles, 2002). The Pe, on the other hand, is a later positive deflection over parietal electrodes, which is assumed to reflect higher-level aspects of error processing that are associated with, or lead to conscious error detection (Endrass, Reuter, & Kathmann, 2007; Overbeek et al., 2005). The Pe has been linked to an accumulation process of evidence that an error has occurred (Murphy, Robertson, Harty, & O’Connell, 2015; M. Steinhauser & Yeung, 2010, 2012) but also to metacognitive concepts such as the graded confidence in a correct decision (Boldt & Yeung, 2015), motivational aspects (Drizinsky, Zülch, Gibbons, & Stahl, 2016; Kim, Marulis, Grammer, Morrison, & Gehring, 2017; Moser, Schroder, Heeter, Moran, & Lee, 2011; Schroder & Moser, 2014), and the negative affective response to the error (Falkenstein, Hoormann, Christ, & Hohnsbein, 2000; van Veen & Carter, 2002).

Based on the shared time course and scalp topographies of the P300 and the Pe (Leuthold & Sommer, 1999; Overbeek et al., 2005), it has been suggested that the stimulus-locked P300 and the response-locked Pe are eventually based on the same neurocognitive processes (Ridderinkhof, Ramautar, & Wijnen, 2009). In fact, both components have been linked to joint activation in bilateral prefrontal and parietal brain regions (Dehaene et al., 1998; Hester, Foxe, Molholm, Shpaner, & Garavan, 2005; Sergent & Dehaene, 2004; Sigman & Dehaene, 2008). This can well explain why particularly the Pe is affected by interference from competing tasks: the Pe interferes with subsequent stimulus processing (Buzzell et al. 2017, but see Beatty et al., 2018) and only the Pe has been shown to be impaired when two temporally overlapping tasks have to be executed (Weißbecker-Klaus et al., 2016).

Whereas the similarity between the Pe and P300 can explain why specifically the Pe suffers from dual-tasking interference, little is known how the brain maintains the ability to reliably detect errors under dual-tasking. One solution to this problem would be to serialize not only task processing but also the accompanying error monitoring processes (Hochman & Meiran, 2005). Processing a T1 error could lead to an additional deferment of T2 processing, an effect that has indeed been demonstrated in behavioral studies (Jentzsch & Dudschig, 2009; M. Steinhauser, Ernst, & Ibald, 2017). However, reduced error signaling rates when an erroneous response is quickly followed by another task (Rabbitt, 2002) and the observation of a reduced Pe in dual-tasking (Weißbecker-Klaus et al., 2016) suggest that such a *serialization* cannot fully protect error monitoring from dual-task interference. Consequently, it might be necessary to actively defer particularly the more resource-consuming aspects of error processing to the end of a dual-task scenario. In this case, processing a T1 error would at least to some degree not occur until the response to T2. Such an *adaptive rescheduling* of conscious error processing would imply that the neural processes that underly the Pe can be detached from early error signals represented by the Ne/ERN.

The present study investigated this adaptive rescheduling hypothesis, which proposes that the brain adapts to dual-task interference in error monitoring by deferring higher-level aspects of error processing to after completion of the dual task. To this end, we focused on T1 errors but analyzed ERPs following both the T1-response and the T2-response. We contrasted trials with a sufficiently long interval between the two task stimuli that allowed serial task execution (1200 ms) with trials that required overlapping task execution due to a considerably shorter interval (300 ms). In serial task execution, we expected to see the typical Ne/ERN and Pe after the erroneous T1-response. Crucially, in overlapping task execution, we predicted that the Pe after the T1-response would be reduced or even absent while a Pe would now be obtained after the correct T2-response. Such a result would demonstrate that higher-level aspects of T1 error processing are deferred at least partially until the completion of T2, and extend previous accounts on an active deferment of component processes in dual-tasking (Hesselmann et al., 2011; Sigman & Dehaene, 2008; R. Steinhauser & Steinhauser, 2018) to the area of performance monitoring.

## Results

### Experiment 1

24 healthy adult participants worked on a variant of the psychological refractory period (PRP) paradigm, which is commonly used to investigate mutual interference between subtasks in dual-tasking situations (Pashler, 1994; Tombu & Jolicœur, 2003). The details of the experimental paradigm can be found in Figure 1.

**Figure 1.**
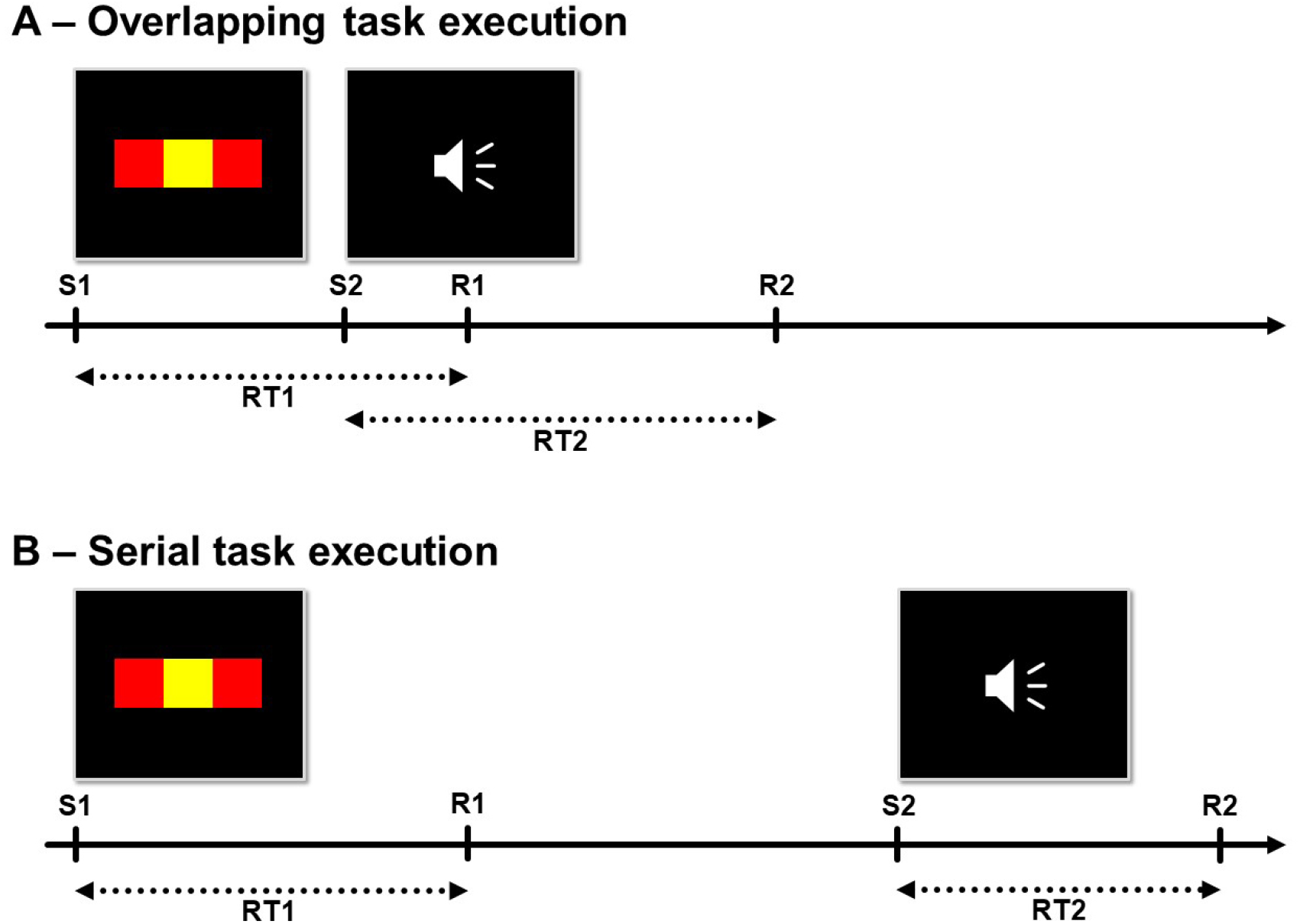
Time course of two example trials. On each trial, participants had to respond to two tasks that were separated by an unpredictable stimulus onset asynchrony of 300 ms (overlapping task execution, A) or 1200 ms (serial task execution, B). A three-choice color variant of the flanker task, in which the color of the central square had to be indicated (red, yellow, blue), served as Task 1. A two-choice pitch discrimination task was presented subsequently as Task 2, with an auditory sine tone stimulus of 400 Hz or 900 Hz. Participants were instructed to respond as fast as possible to both tasks, in the given order. S1 = Stimulus 1. S2 = Stimulus 2. R1 = Response 1. R2 = Response 2. RT1 = response time to Task 1. RT2 = response time to Task 2.

### Behavior

RTs and error rates are depicted in Figure 2. To verify that our paradigm creates a dual-tasking scenario with overlapping task execution, we first examined whether two typical effects of dual-tasking can be observed in this dataset: First, the so-called PRP effect refers to the observation that RTs to T2 increase with a decreasing stimulus onset asynchrony (SOA), and thus indicates a form of dual-task cost. This effect is typically explained by the idea that, with more overlap between tasks, T2 execution is delayed (Pashler, 1994) or suffers from depleted resources (Tombu and Jolicœur, 2003). Second, it has recently been shown that T1 errors lead to increased RTs to T2, and this post-error slowing is larger for short SOAs than for long SOAs (Steinhauser et al., 2017). This phenomenon presumably reflects interference between T1 error monitoring and execution of T2, which again is larger with short SOAs.

**Figure 2.**
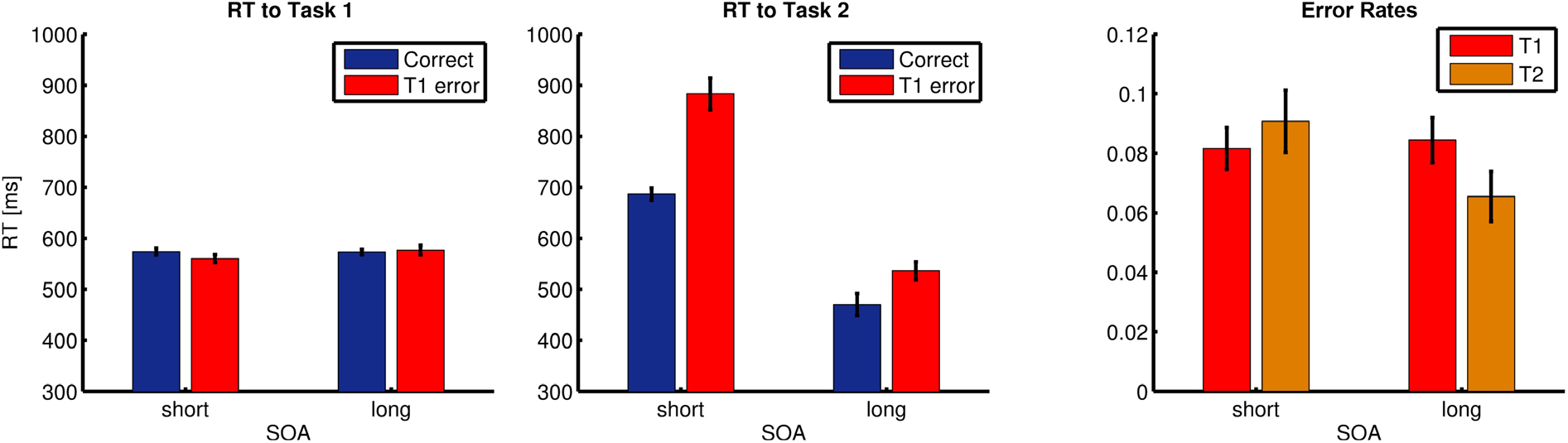
Behavioral results of Experiment 1. RTs to Task 1 and Task 2 are depicted in the left and middle panel, respectively. Error rates are presented in the right panel. Error bars indicate within-subject standard errors of the mean (Cousineau, 2005; Morey, 2008). RT = response time. SOA = stimulus onset asynchrony. T1 = Task 1. T2 = Task 2.

To analyze both effects in our data, we subjected RTs for T1 and T2 separately to repeated measures ANOVAs on the variables SOA (short vs. long) and T1 Correctness (correct vs. error). Characteristic of PRP paradigms, no significant effects were found for RTs to T1, *F*s(1, 23) < 2.70, *p*s > .11, *η*_*p*_^2^ < .11. In contrast, RTs to T2 were considerably slower on trials with short SOA than with long SOA, *F*(1, 23) = 83.4, *p* < .001, *η*_*p*_^2^ = .78, reflecting the typical PRP effect. In addition, RTs to T2 were slower when the preceding T1-response was an error, *F*(1,23) = 56.8, *p* < .001, *η*_*p*_^2^ = .71, and as expected, this post-error slowing on T2 was increased with short SOA, *F*(1, 23) = 25.9, *p* < .01, *ηp*² = .53. Thus, our data replicate well-known signatures of dual-task interference, which demonstrates that the short-SOA condition induced overlapping task execution. In addition, an analysis of inter-response intervals (IRIs, the time between the T1 response and the T2 response) in the short-SOA condition ruled out that participants had grouped their responses. While IRIs were considerably longer on T1 errors (587 ms) compared to corrects (381 ms) due to the above mentioned post-error slowing, *t*(23) = 8.35, *p* < .001, *d* = 1.25, IRIs in both conditions were far beyond the commonly used threshold for response grouping (i.e., simultaneously pressing both response buttons) of 50 – 100 ms (Hommel, 1998; Pashler & Johnston, 1989; Welford, 1952).

Error rates for T1 were high enough for an analysis of error-related brain activity. A mean T1 error rate of 8.31 % resulted in an average number of 98.3 trials with T1 errors but correct T2 responses per participant. A repeated measures ANOVA on the variables SOA (short vs. long) and Task (T1 vs. T2) yielded a significant interaction, *F*(1, 23) = 8.06, *p* = .009, *η*_*p*_^2^ = .26, indicating that, whereas T1 errors were equally frequent in short-SOA and long-SOA trials, T2 errors were more frequent in short-SOA trials as compared to long-SOA trials, *t*(23) = 2.69, *p* = .013, *d* = .50. This again demonstrates increased interference in the short-SOA condition.

### Ne/ERN and Pe

To investigate our rescheduling hypothesis, we analyzed error-related brain activity in short-SOA trials, which induce overlapping task execution, and long-SOA trials, which allow for serial task execution. Our central prediction was that, with a short SOA, the Pe associated with T1 errors should partially be rescheduled to the end of the dual-task. This would result in a reduced Pe following incorrect T1 responses but the emergence of a Pe after correct T2 responses. We had no explicit hypothesis on the Ne/ERN but nevertheless analyzed this component to determine whether comparable effects can be found for this early form of error processing.

We initially analyzed T1-response-locked ERPs to find out about immediate neural correlates of error processing after T1 errors. A distinct parietal positivity, the Pe, was clearly observable for both SOA conditions (Fig 3A). For an analysis of the Pe, we subjected mean amplitudes at electrode POz to a repeated measures ANOVA on the variables SOA (short vs. long) and T1 Correctness (correct vs. error). A Pe was evident across both SOA conditions, as indicated by a main effect of T1 Correctness, *F*(1, 23) = 28.6, *p* < .001, *η*_*p*_^2^ = .55, but a significant interaction revealed that this Pe was far less pronounced in trials with short SOA than in trials with long SOA, *F*(1, 23) = 11.0, *p* = .003, *η*_*p*_^2^ = .32. Raw ERP waves in Figure 3 suggest that this interaction is mainly driven by a difference in the amplitudes of correct trials. This is likely caused by the fact that in most trials of the short SOA condition, the stimulus of T2 is presented and processed sometime within the time window observed here, eliciting stimulus-locked ERPs that are superimposed on the raw ERP waveforms of both correct and error trials in the short SOA condition (see Sigman & Dehaene, 2008). For this reason, we quantify the Pe not from error trials alone but as the difference between correct and error trials. To additionally rule out that the observed interaction may truly be rooted in differences in correct trials, we correlated the difference wave of conditions correct long and correct short with the difference wave of the interaction term of the above analysis. We found no significant correlation, *r* = 0.14, *p* = .50, indicating that the Pe reduction in trials with short SOA is not a mere consequence of the amplitude difference in correct trials.

**Figure 3.** ERPs locked to the T1-response at posterior (A) and frontocentral (B) electrodes. Difference waves are computed from the respective T1 error minus T1 correct raw ERPs. Scalp topographies represent these difference waves. Gray areas indicate the time intervals for statistical testing for the Pe (A) and the Ne/ERN (B). T1 = Task 1. SOA = stimulus onset asynchrony.

Figure 3B shows that also a clear frontocentral Ne/ERN was observable in both SOA conditions. Indeed, subjecting mean amplitudes at electrode FCz to a repeated measures ANOVA of the same variables as above revealed that a significant Ne/ERN across both SOAs was obtained, *F*(1, 23) = 22.8, *p* < .001, *η*_*p*_^2^ = .50. In addition, a main effect of SOA demonstrated that mean amplitudes of correct as well as error trials were more negative in trials with short SOA, *F*(1, 23) = 13.5, *p* = .001, *η*_*p*_^2^ = .37. However, the interaction between both variables did not reach significance, *F*(1, 23) = 2.63, *p* = .12, *η*_*p*_^2^ = .10.

Having shown that higher-level aspects of error processing as reflected by the Pe were largely impaired in the short-SOA condition, we subsequently examined neural correlates of T1 error processing after completion of the whole dual-task, i.e., in T2-response-locked data (Fig. 4). We investigated the possible emergence of such a deferred Pe (Fig. 4A) by means of ANOVAs on the variables SOA (short vs. long) and T1 Correctness (T1 correct vs. T1 error). It must be noted that both conditions of the variable T1 Correctness here represent trials whose T2 was answered correctly, i.e., the “error-related” brain activity reported here was locked to a correct T2-response. Nonetheless, we obtained a significant interaction between SOA and T1 Correctness on Pe amplitudes, *F*(1, 23) = 5.55, *p* = .027, *η*_*p*_^2^ = .19, reflecting that correct T2-responses elicited a significant Pe on short-SOA trials, *t*(23) = 2.00, *p* = .029, *d* = .27, but not on long-SOA trials.

**Figure 4.** ERPs locked to the (correct) T2-response at posterior (A) and frontocentral (B) electrodes. Difference waves are computed from the respective T1 error minus T1 correct raw ERPs. Scalp topographies represent these difference waves. Gray areas indicate the time intervals for statistical testing for the Pe (A) and the Ne/ERN (B). T1 = Task 1. SOA = stimulus onset asynchrony.

The same analysis on the Ne/ERN amplitudes revealed only a marginally significant main effect of T1 Correctness, *F*(1, 23) = 3.48, *p* = .075, *η*^2^_part._ = .13, indicating a negativity after T1 errors compared to T1 correct trials, but no significant interaction, *F*(1, 23) = 1.73, *p* = .20. Visual inspection of the topographies (Fig. 4B) revealed that this effect peaked earlier than the typical Ne/ERN and had a more frontal distribution than the Ne/ERN in T1-response-locked potentials (however, the same analysis at electrode Fz showed similar results). This suggests that, while there appears to be some frontocentral activity related to the T1 error in T2-response-locked data, this effect lacks robustness and differs from the usually observed Ne/ERN.

To sum up, we could provide evidence that overlapping task execution goes along with a reduction of the Pe to the incorrect T1 response whereas a sizeable Pe emerges after the correct T2-response. We interpret this *deferred Pe* as a rescheduling of higher-level error processing to the end of the dual-task trial.

### Relationship between immediate and deferred error processing

Although the immediate Pe that followed the T1 response was strongly reduced in the short-SOA condition, it was observable there nonetheless. This suggests two possible ways of how higher-level error monitoring is carried out under conditions of overlapping task execution. On the one hand, error monitoring after the T2 response could occur predominantly on those trials that also feature strong error monitoring immediately after the T1 response. An actual rescheduling mechanism as outlined above, which defers T1 error processing to after completion of T2, however, would by definition require the exact opposite pattern: error monitoring after the T2 response would have to occur on error trials that did not exhibit considerable immediate error processing after T1. To distinguish between these two possibilities, we examined the trial-wise relationship between immediate and deferred Pe. This was done in two methodologically different ways, to combine their respective advantages and countervail their disadvantages.

First, we investigated the inverse relationship of immediate and deferred error processing by deriving single-trial estimates for the R1-locked and the R2-locked Pe from mean amplitudes in raw data and comparing them in a regression-based analysis. This analysis had to be limited to trials with IRI > 400 ms, however, because raw EEG data within the same epoch are highly susceptible to autocorrelations in overlapping time periods. Importantly, this does not affect the remaining analyses of the present study, because there, EEG data from different, individually baseline-corrected epochs are compared. The restriction to IRIs above 400 ms resulted in M = 31.04 trials per participant. Figure 5 depicts strong negative beta weights in a significant cluster from 290 ms onwards in R1-locked data and 170 ms in R2-locked data, that is, around the time of the R1-locked and R2-locked Pe. This is evidence for a trade-off between the immediate and the delayed Pe, supporting the account that error processing happens immediately after some erroneous responses, whereas it is deferred on other trials.

**Figure 5.** Regression weights of the regression-based single-trial analyses for Experiments 1 and 2, in which amplitudes of R1-locked data predict amplitudes of R2-locked data. Black lines indicate the borders of significant clusters as revealed by a cluster-based permutation test. Red dotted lines indicate the peak amplitude of the Pe in the respective difference waves. A significant negative association of the R1-locked Pe and R2-locked Pe becomes evident in both experiments.

Second, we used multivariate pattern analysis (MVPA) to create a set of classifiers over consecutive time windows that optimally distinguished between correct trials and T1 error trials based on T1-response-locked brain activity. This approach allowed us to examine all trials irrespective of their IRI and is furthermore less susceptible to spurious non-phase-locked activity in the EEG data (Parra et al., 2002; Parra, Spence, Gerson, & Sajda, 2005). Mean classifier accuracy peaked around 190 ms with an Az of .61 but inspection of participants’ individual classifier accuracies revealed that one participant (Subject 13) exhibited a remarkably low classification accuracy of .34, which is more than 2.5 standard deviations below the mean classification accuracy across all participants. This apparently failed attempt to compute a successful MVPA for this participant is likely due to the small number of 20 trials that entered the training set (the average training set size of all participants was 120.63 trials). For this reason, data from Subject 13 was excluded from subsequent MVPA-based analyses. Nonetheless, all effects remain significant also when including that participant (all *p*s < .05). Based on the peak classifier window, we computed the prediction value for each trial. These prediction values represent the degree to which each trial elicits error-related brain activity in this time window (Boldt & Yeung, 2015; Steinhauser & Yeung, 2010), and thus represent a single-trial estimate of the T1-response-locked Pe. We utilized these prediction values to split the error trials of each participant into three equally-sized bins that represent a small, medium, and large T1-response-locked Pe (Fig. 5A). A one-way repeated measures ANOVA on mean Pe amplitudes with the variable Bin (small Pe vs. medium Pe vs. large Pe) showed that this MVPA-based separation process was in fact able to separate trials according to Pe size, *F*(2, 44) = 4.72, *p* = .015, *η*_*p*_^2^ = .18. Contrasts revealed that this effect was mainly driven by trials in the large Pe bin having a larger T1-response-locked Pe than trials in the small Pe bin, *t*(22) = 2.90, *p* = .008, *d* = .52. Crucially, an equivalent analysis of T2-response-locked Pe values based on this T1-response-based bin separation (Fig. 5B) shows the opposite pattern, *F*(2, 44) = 6.75, *p* = .004, *η*_*p*_^2^ = .23. Trials with a large immediate Pe following the T1-response showed a particularly small deferred Pe following the T2-response, *t*(22) = 4.10, *p* < .001, *d* = .63. This conceptually replicates the above regression-based findings on a trade-off between the immediate and deferred Pe and thus provides additional support for the idea that the rescheduling of higher-level error processing occurs predominantly on trials with little immediate error processing. This finding additionally rules out the above-mentioned alternative interpretation that differences in the T1-response-locked Pe with regard to short and long SOA could be rooted only in differences between the correct trials in the two conditions. MVPA-based decoding of the degree of immediate error processing directly modulates Pe amplitudes of error trials as well (Fig. 5A).

The MVPA-based differentiation of trials with small, medium and large immediate Pe eventually also allowed us to investigate how immediate error processing affects selecting and executing the response to Task 2 (Fig. 5C). We compared IRIs in a one-way repeated measures ANOVA (small vs. medium vs. large immediate Pe) and found a significant difference between the conditions, *F*(2,44) = 3.42, *p* = .045, *η*_*p*_^2^ = .13. Contrasts confirmed that trials with a large immediate Pe were followed by slower responses to T2 than trials with a small immediate Pe, *t*(22) = 2.94, *p* = .001, *d* = .40, which in turn supports the idea that immediate error processing negatively affects efficient execution of the second task.

### Experiment 2

Individual trials that elicit a deferred Pe with simultaneous omission of an immediate Pe could also be rooted in the problem of credit assignment (Fu & Anderson, 2008; Sutton & Barto, 1998; Walsh & Anderson, 2011). It is conceivable that trials with a deferred Pe actually reflect that error monitoring occasionally misattributed internal error signals to T2, and therefore falsely detected a T2 error. Rather than a rescheduling of T1-error processing, such a Pe would represent immediate T2-error processing. In a follow-up experiment with 24 different participants (out of which one participant had to be excluded due to technical difficulties during data acquisition), we did not only want to replicate our initial findings but also investigated whether this alternative explanation could account for our findings.

In Experiment 2, participants worked on the same PRP paradigm but reported after each trial by key press, whether they had committed an error in T1, T2, or both. This allowed us to distinguish between correctly assigned T1 errors (T1 hits) and T1 errors that were mistakenly reported as T2 errors (T2 false alarms). Rates of averaged error reports are depicted in Figure 7A. In fact, only 4.9% of all T1 errors were T2 false alarms (on average 2.1 trials per participant) and thus occurred very rarely. As only 17.0% of T1 errors were reported to be correct (T1 misses; on average 6.78 trials per participant), we could base our further ERP analyses on a solid subset of 73.8% of T1 hits (on average 31.0 trials per participant).

To compensate for the increased trial duration due to trial-wise error reporting, Experiment 2 was restricted to the short SOA condition. Only T2-response-locked data are reported here as no immediate Ne/ERN and Pe could be compared between SOA conditions. Although only T1 hits were included in the analysis, a clear deferred Pe could be observed after T2-response-locked data (Fig. 6B), *t*(22) = 2.86, *p* = .009, *d* = .57.

**Figure 6.** ERPs at a posterior electrode locked to the T1-response (A) and T2-response (B) and inter-response intervals after T1 errors (C). T1 error trials are divided into three separate conditions – small, medium, and large immediate Pe – based on a single-trial estimate of the Pe. Correct trials are presented for comparison (thin line). A 20 Hz lowpass filter was applied for better visibility. Gray areas indicate the time intervals for statistical testing for the Pe. T1 = Task 1. T2 = Task 2.

As in Experiment 1, the analysis of the Ne/ERN in T2-response-locked data revealed only a marginally significant negativity for T1 errors relative to correct trials at electrode FCz, *t*(22) = 1.87, *p* = .075, *d* = .51. However, inspection of the topographies (Fig. 7B) showed that the peak of the Ne/ERN was again at more frontal electrodes. Correspondingly, additional testing was conducted at electrode Fz and now revealed a significant deferred Ne/ERN, *t*(22) = 2.26, *p* = .034, *d* = .62.

**Figure 7.**
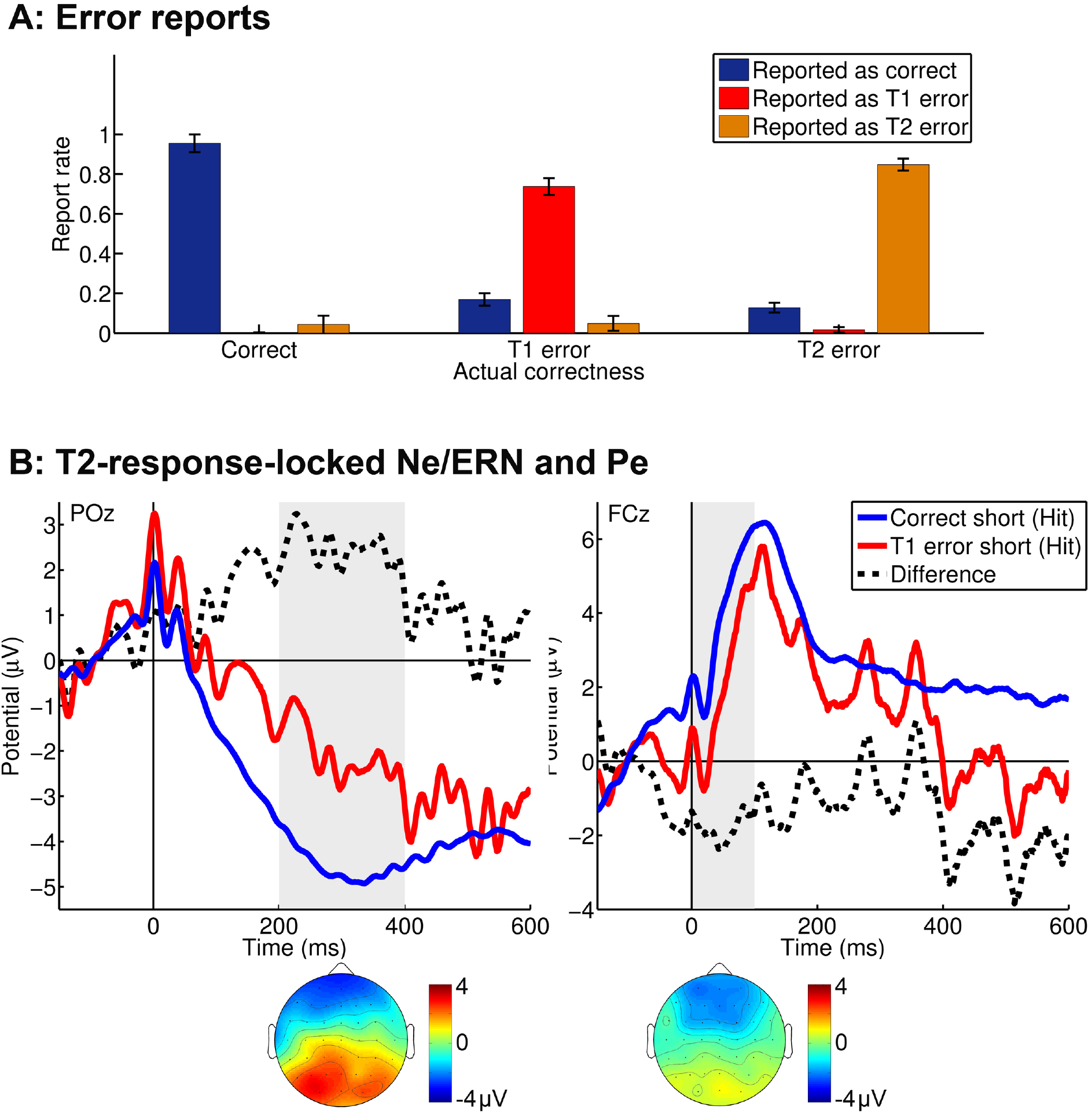
Error reports (A) and ERP results (B) of Experiment 2. Error bars in Panel A indicate within-subject standard errors of the mean (Cousineau, 2005; Morey, 2008). Panel B represents ERPs locked to the T2-response at posterior (left) and frontocentral (right) electrodes. Difference waves (dotted lines) are computed from T1 error minus correct raw ERPs. Scalp topographies represent these difference waves. Gray areas indicate the time intervals for statistical testing for the Pe (left) and the Ne/ERN (right). T1 = Task 1. T2 = Task 2.

The regression-based approach on raw data with IRI > 400ms yielded a pattern with close similarity to that in Experiment 1, albeit somewhat earlier in R2-locked data (Figure 5). Again, strong negative beta weights form a significant cluster from 290 ms onwards in R1-locked data and between 125 ms and 230 ms in R2-locked data. We tentatively suggest that this negative correlation between the R1-locked Pe and the R2-locked Pe is reduced in the R2-locked time window beyond 250 ms due to the strong influence of a pronounced overlaying Contingent Negative Variation in anticipation of the error report (see Figure 7B). A robust MVPA-based bin separation of trials with large vs. medium vs. small immediate Pe as in Experiment 1 was not possible for Experiment 2 because classifier training did not yield a classification accuracy beyond the 5% significance threshold as established by a permutation test, likely due to the smaller size of the training data set.

Taken together, we could replicate the deferred Pe in Experiment 2. As this analysis was restricted to those error trials that were correctly reported as T1 errors, we can rule out that this deferred Pe originates in occasional T1-error trials that are misattributed as T2-errors.

## Discussion

In two ERP experiments, we investigated neural correlates of error monitoring in dual-tasking. Previous studies suggested that increased interference due to temporally overlapping tasks is countered by an active deferment of task components (Hesselmann et al., 2011; Meyer & Kieras, 1997; Sigman & Dehaene, 2008). Based on this idea, we hypothesized that because higher-level processing of T1 errors can interfere with the execution of T2 (M. Steinhauser et al. 2017), also performance monitoring, more precisely the more resource-consuming, higher-level aspects of T1 error processing, are adaptively rescheduled to after the completion of the whole dual-task. To this end, we analyzed the Pe, an ERP component that is commonly reported to represent such higher-level, central processing of the error and involves aspects such as evidence accumulation (Murphy et al., 2015; M. Steinhauser & Yeung, 2010, 2012), decision confidence (Boldt & Yeung, 2015), motivation (Drizinsky et al., 2016; Kim et al., 2017; Moser et al., 2011; Schroder, Moran, Donnellan, & Moser, 2014), and affect (Falkenstein et al., 2000; van Veen & Carter, 2002). In both experiments, we found a result pattern that confirms our prediction. When the execution of two tasks overlaps, errors in the first task lead to a reduced immediate Pe whereas a deferred Pe appears after the response to the second task.

Our results are in line with the idea that higher-level error processing as reflected by the Pe interferes with the execution of a subsequent task (Buzzell et al., 2017; Weißbecker-Klaus et al., 2016). In our study, only the Pe but not the Ne/ERN after T1 errors was reduced when task execution was overlapping. This conforms with recent findings by Weißbecker-Klaus et al. (2016), who found a similar reduction of the Pe when a flanker task was executed concurrently with a semantic task. In both studies, this impairment to immediate error detection likely originates from interference by T2, which requires cognitive resources for T2-response selection at the same time as T1-error processing would occur. The decision process for selecting the correct response to T2 was previously suggested to rely on similar resources as error processing (Hochman & Meiran, 2005). On the neural level, this can well be explained by recent electrophysiological studies that view also higher-level error processing as a decision process (Wessel, 2012; Wessel, Danielmeier, & Ullsperger, 2011) and studies that highlight the physiological similarities of the P300 and the Pe (Leuthold & Sommer, 1999; Overbeek et al., 2005). Given that interference between central decision processes is seen as the main origin of dual-task costs (Pashler 1994; Tombu and Jolicœur, 2003), this explains why particularly the Pe as a neural correlate of higher-level error detection (Overbeek et al., 2005) is impaired in dual-tasking.

In addition to a reduced immediate Pe after the erroneous T1 response, we found a deferred Pe after completion of the dual-task even though the second response itself was correct. MVPA-based single-trial estimates of the Pe as well as a regression-based approach on raw data suggest that the deferred Pe occurred predominantly on trials with little immediate error processing. This temporal detachment of higher-level error processing from the early error signals that are represented by the Ne/ERN – on average by more than 580 ms – goes beyond a mere serialization as it was previously found for the P300 in response selection (Hesselmann et al., 2011; Marti et al., 2015; Sigman & Dehaene, 2008). In the present study, at least some aspects of higher-level error detection appear to be suspended until a whole additional task has been executed. Analysis of IRIs furthermore confirmed that the deferment of error processing is associated with faster responses to T2, suggesting reduced dual-tasking interference on such trials, in line with Buzzell et al.’s (2017) finding that a smaller Pe is also linked to improved sensory processing in the subsequent trial. This is strong support for the adaptive rescheduling account of error detection in dual-tasking, which suggests such a mechanism to reduce interference between the subtask representations and hence to maintain the ability to detect and evaluate errors. Notably, time course and scalp topography of the deferred Pe indicate that this phenomenon affects what has previously been described as the posterior “late Pe”, not the earlier frontocentral aspects of the Pe (Endrass et al., 2007; Ullsperger et al., 2014, see also Steinhauser & Yeung, 2010).

Previous research on error detection as an evidence accumulation process (Murphy et al., 2015; M. Steinhauser & Yeung, 2010, 2012) may help to approach the neural basis of this deferment mechanism. The evidence accumulation account considers error detection as a decision process (M. Steinhauser & Yeung, 2010) and is thus well compatible with the serialization of central response selection processes in dual-tasking (Pashler, 1994; Tombu & Jolicœur, 2003). The idea that the Pe represents the ongoing accumulation of evidence that an error has occurred by mirroring the internal weights of evidence can consequently be linked to the processes that form the basis of the stimulus-locked P300 (see also Ridderinkhof et al., 2009). Marti, King, and Dehaene (2015) recently showed that the brain networks linked to the emergence of the P300 bring about a competition between the neural representations of the subtasks in dual-tasking, so that only one of those representations can be active at a time and a deferment of the T2-related P300 is induced (see also Hesselmann et al., 2011; Sigman & Dehaene, 2008). Shared neural generators of the Pe and the P300 could therefore also lead to a deferment of the evidence accumulation process for the T1 error until completion of T2, thus constituting the deferred Pe that was observed in the present two experiments.

Analyzing brain activity in a PRP paradigm bears the risk that activity related to T1 is accidentally attributed to T2 (Sigman & Dehaene, 2008). For instance, what appears to be a deferred Pe after the T2-response could simply be a carry-over of the immediate Pe after the T1-response. However, there are several reasons why this cannot account for the present results. First, the deferred Pe shows a distinct time course, emerging about 200 ms after the T2 response, which is comparable to the latency of the immediate Pe as well as of the Pe in other studies (Overbeek et al., 2005). Second, even with a short SOA of 300 ms, the average interval between the T1-response and T2-response on error trials was 587 ms. That is, the onset of the immediate Pe is clearly prior to the T2-response, and any sustained differences between T1 errors and correct trials emerging prior to the T2-response are controlled by the pre-response baseline. Finally, and most importantly, our results indicate an inverse correlation between immediate Pe and deferred Pe. If the deferred Pe reflected a carry-over of activity from the immediate Pe, both effects should be positively correlated.

In Experiment 2, we could also rule out that the deferred Pe results from internal misattribution of occasional T1 errors as T2 errors, which would imply a credit assignment problem (Fu & Anderson, 2008; Sutton & Barto, 1998; Walsh & Anderson, 2011) as the origin of our findings. Trial-wise error-reports in Experiments 2 revealed that credit-assignment errors occurred extremely rarely in our tasks. Moreover, we obtained a deferred Pe even on the subset of correctly assigned T1 errors. This suggests that even when higher-level processing of the T1 error is postponed, these errors are still detected and correctly assigned to their corresponding task. One could speculate that the proposed scheduling mechanism even contributes to solving the credit assignment problem. On the one hand, rescheduling counteracts interference between T1 error processing and T2 response selection which otherwise would increase the risk of confusing error signals from T1 and T2 responses. On the other hand, when higher-level error evaluation occurs after both responses were executed, the involved decision process could utilize cues from both tasks to correctly assign error signals (like post-response conflict) to their corresponding tasks.

Though far less robust than the deferred Pe, we also obtained frontocentral activity related to T1 errors following the T2 response. This could reflect a deferred Ne/ERN, although this activity had a slightly different time course and spatial distribution as the immediate Ne/ERN after T1-responses. However, this deferred Ne/ERN differed in a crucial aspect from the deferred Pe as it was not associated with a reduced immediate Ne/ERN. Without this trade-off between immediate and deferred error processing, this effect cannot reflect a scheduling mechanism. It is also implausible that the deferred Ne/ERN reflects post-response conflict induced by a corrective response tendency (Yeung et al., 2004) or a reward prediction error (Holroyd & Coles, 2002) as both these mechanisms are temporally linked to the erroneous response. Instead, the deferred Ne/ERN could represent a negative affective signal to the T2-response. Several studies could show that the Ne/ERN involves early aspects of affective processing of an error (Aarts, Houwer, & Pourtois, 2013; Maier, Scarpazza, Starita, Filogamo, & Ladavas, 2016). The deferred Ne/ERN hence could reflect that committing an error in one task leads to a devaluation also of the second task, indicating that the whole dual-task trial is processed as erroneous.

Taken together, our results suggest that reliable error detection in dual-tasking is maintained by a mechanism that adaptively reschedules higher-level aspects of error processing to after the completion of the whole dual-task. Our findings indicate that the flexible reorganization of componential processes under dual-tasking is not only a viable strategy to prevent interference between decision processes involved in task execution (Hesselmann et al., 2011; Marti et al., 2015; Meyer & Kieras, 1997; Sigman & Dehaene, 2008), but also serves to reduce interference between task execution and performance monitoring.

## Materials and methods

### Participants

We conducted two separate, consecutive experiments with different groups of participants. 24 healthy students (2 male; age: M = 22.3 years; SD = 4.5 years; 2 left-handed) participated in Experiment 1, and 24 healthy students (2 male; age: M = 24.0 years; SD = 3.9; 2 left-handed) participated in Experiment 2. With an assumed correlation of .5 among cell means, a sample size of N = 24 should allow for detecting medium-sized effects (*f* = .25) in the present repeated-measures design with a statistical power of .82. All participants were recruited from the Catholic University of Eichstätt-Ingolstadt and received course credit or payment (8€ per hour). Informed consent was provided by all participants and the study was approved by the ethics committee of the Catholic University of Eichstätt-Ingolstadt. One subject in Experiment 2 had to be excluded from further analysis due to technical problems during data acquisition.

### Task and procedure

#### Experiment 1

Adopting the PRP paradigm from Steinhauser, Ernst, and Ibald (2017), we combined an error-prone three-choice color flanker task and a two-choice pitch discrimination task. The flanker task stimulus (see Fig. 1) consisted of three horizontally arranged squares of .82° edge length. The central target square and the two flanker squares were either red, yellow or blue. While both flanker squares had the same color, this color was always different from that of the target. As three colors were used, it was possible to exclusively present these more error-prone incongruent stimuli without enabling participants to infer the target color from the flankers. Stimuli for the secondary pitch task were low (400 Hz) and high (900 Hz) sine tones.

The trial procedure is depicted in Figure 1. Participants first had to respond to the flanker task (T1) and then to the pitch discrimination task (T2). Each trial started with the presentation of a fixation cross for 500 ms. Following this, the flanker task was presented by initially displaying the two flankers for 60 ms and then the target together with the flankers for 200 ms. The pitch discrimination task started after an SOA of either 300 ms (short) or 1200 ms (long), which was chosen randomly within the blocks. The stimulus for the pitch task was presented for 150 ms. After responses to both tasks were given, the fixation cross of the next trial was presented 500 ms after the response. The short SOA was supposed to create a dual-tasking situation with overlapping task execution while at the same time preventing excessive response grouping, which was observed in a piloting phase with even shorter SOAs. In contrast, the long SOA was intended to serve as a baseline that allows for serial task execution.

For the flanker task, participants had to indicate the color of the central target square by pressing the “Y”, “X”, or “C” button on a German QWERTZ keyboard (in which Y and Z are exchanged) with their left hand (the T1-response). For the pitch discrimination task, participants entered their response with “arrow down” for a low pitch and “arrow up” for a high pitch (the T2-response). The mapping of colors to keys was counterbalanced across participants, and for one half of the participants, the task-to-hand assignment was reversed (resulting in “,”, “.”, “-” for the flanker task and “A” and “Y” for the pitch task). If the T2-response was given before the onset of the pitch stimulus, written feedback (“AUFGABE 2 ZU FRÜH!”, *engl*. task 2 too early) was given immediately and the trial was excluded from data analysis.

The experiment started with a series of practice blocks that ensured that participants had thoroughly learned the experimental paradigm and the mappings of colors/pitches to keys. First, in three blocks of 24 trials the flanker task was presented alone. Then, one block of 16 trials served to practice the pitch task alone, and in one subsequent block of 36 trials, both tasks were responded to together. Actual testing consisted of 10 blocks of 108 trials each, resulting in a total number of 1080 trials. Oral feedback to respond faster was given after blocks, if the error rate for any of the tasks was below 10%.

#### Experiment 2

Experiment 2 adopted the experimental method of Experiment 1 but added a trial-wise report of errors. To this end, response keys to the two tasks were changed (“A”, “S”, “D” and “L”, “P” for one half of participants, “L”, “Ö”, “Ä” and “A”, “Q” for the other half) and two additional keys, “alt” and “alt gr”, served to report errors. In each trial, 600 ms after the response to Task 2, an additional instruction (“FEHLER?”, *engl*. error?) appeared for 1000 ms in the center of the screen. During this time, participants should indicate if they had just committed an error in the task of their left hand (“ALT”) or their right hand (“ALT GR”), or both. No response was required if the participants considered both their responses correct. To account for the increased trial duration, Experiment 2 was restricted to only the short SOA of 300 ms. With 10 blocks of 84 trials each, this resulted in 840 trials for the condition with overlapping task execution.

### Data acquisition

The EEG was recorded from 64 electrodes at a sample rate of 512 Hz with a BioSemi Active-Two system (BioSemi, Amsterdam, The Netherlands; channels Fp1, AF7, AF3, F1, F3, F5, F7, FT7, FC5, FC3, FC1, C1, C3, C5, T7, TP7, CP5, CP3, CP1, P1, P3, P5, P7, P9, PO7, PO3, O1, Iz, Oz, POz, Pz, CPz, Fpz, Fp2, AF8, AF4, AFz, Fz, F2, F4, F6, F8, FT8, FC6, FC4, FC2, FCz, Cz, C2, C4, C6, T8, TP8, CP6, CP4, CP2, P2, P4, P6, P8, P10, PO8, PO4, O2 as well as the left and right mastoid). The CMS (Common Mode Sense) and DRL (Driven Right Leg) electrodes were used as reference and ground electrodes. Vertical and horizontal electrooculogram (EOG) was recorded from electrodes above and below the right eye and on the outer canthi of both eyes. All electrodes were off-line re-referenced to averaged mastoids.

### Experimental design and statistical analysis

#### Design

Each trial was assigned to a condition based on the SOA (short, long) and T1 Correctness (correct, error). T1 Correctness relied on the post-hoc classification of whether the response in the flanker task (T1) was correct or not. Trials on which an error in the pitch discrimination task (T2) occurred were removed from all analyses of response times (RTs) and ERP data.

#### Data analysis

For the analysis of RTs, trials were excluded whose RT deviated more than three standard deviations from the RT mean of each condition and participant. Error rates were arcsine-transformed prior to statistical testing (Winer, Brown, & Michels, 1991). All analyses were performed using custom MATLAB v8.2 (The Mathworks, Natick, MA) scripts and EEGLAB v12.0 (Delorme & Makeig, 2004) functions. All data and analysis scripts are publicly available in an online repository hosted by the Open Science Framework (https://osf.io/5ub8z/). For both Experiments, continuous EEG data was initially band-pass filtered to exclude frequencies below 0.1 Hz and above 40 Hz. Then, epochs were created for two separate analyses, first from −500 ms before to 1000 ms after the T1-response (T1-response-locked dataset) and, second, from −500 ms before to 1000 ms after the T2-response (T2-response-locked dataset). Epochs in both analyses were baseline-corrected by subtracting mean activity between −150 ms and −50 ms before the response, as neural correlates of performance monitoring were previously found to emerge slightly before the response button press. Separately for each dataset, electrodes were interpolated using spherical spline interpolation if they met the joint probability criterion (threshold 5) as well as the kurtosis criterion (threshold 5) in EEGLAB’s channel rejection routine (pop_rejchan.m). Epochs were removed that contained activity exceeding +/−300 V in any channel except AF1, Fp1, Fpz, Fp2, AF8 (to prevent exclusion of blink artifacts, which were corrected at a later stage) and whose joint probability deviated more than 5 standard deviations from the epoch mean. To correct for eye blinks and muscular artefacts, an infomax-based ICA (Bell & Sejnowski, 1995) was computed and components with time courses and topographies typical for these artefacts were removed after visual inspection.

In both experiments and for each dataset (T1-response-locked and T2-response-locked), following a considerable number of previous studies that feature a Pe peaking over parieto-occipital electrodes (Beatty et al., 2018; Endrass et al., 2007; Shalgi, Barkan, & Deouell, 2009; M. Steinhauser & Yeung, 2010, 2012), the Pe was quantified by comparing mean amplitudes from 200 ms to 400 ms after the respective response at electrode POz between T1 error trials and correct trials. The Ne/ERN was quantified by comparing mean amplitudes from 0 ms to 100 ms after the respective response at electrode FCz between T1-error trials and correct trials. In Experiment 2, these analyses were restricted to T1-error trials that were reported as T1 errors, and correct trials that were reported as correct trials.

As we found that T1 errors elicited both an immediate Pe in T1-response-locked data and a deferred Pe in T2-response-locked data, we aimed to investigate the relationship between both types of Pe on a single-trial level. Our approach was to categorize trials according to the size of the immediate Pe and then analyze differences between these trials with respect to the deferred Pe. To acquire a robust single-trial estimate of the T1-response-locked Pe, we used a multivariate pattern analysis (MVPA) based on the linear integration method introduced by Parra et al. (2002, 2005). This method has previously been used to quantify error-related brain activity on a single-trial level (Boldt & Yeung, 2015; M. Steinhauser & Yeung, 2010). Here, we provide only a brief description of this method while details can be found elsewhere (e.g., Steinhauser & Yeung 2010). In a first step, we computed a set of classifiers on T1-response-locked data that discriminated optimally between correct trials and T1 errors. Classifiers were constructed for consecutive, partially overlapping time windows from 0 ms to 400 ms after the T1-response (width 50 ms, step size 10 ms). All classifiers were trained, separately for each participant, on T1 error trials and the same number of randomly drawn correct trials. Then, the classifier was selected that featured the highest discrimination sensitivity, as indicated by the Az score. To prevent overfitting, Az was computed using leave-one-out cross-validation. Using this classifier, we calculated prediction values for each error trial. These prediction values represent single-trial estimates of error-related brain activity in the respective classifier window. Based on these estimates, we assigned each error trial to one of three equally sized bins: small Pe, medium Pe, and large Pe in T1-response-locked data. By analyzing Pe amplitudes in T2-response-locked data according to these bins, we were able to investigate the relationship between immediate and deferred Pe elicited by T1 errors.

As a second way of investigating a possible trade-off between the R1-locked Pe and the R2-locked Pe, we computed an analysis based on linear regression on raw data. To get a sufficiently robust measure of the Pe in single trials, we computed mean amplitudes of trial-wise EEG data in a 100 ms window on a cluster of posterior electrodes (Pz, P1, P2, POz, P03, P04). This was done in steps of 10 ms on time windows centering from 0 ms to 400 ms after both responses. For each such combination of single-trial Pe values in R1-locked data and R2-locked data, the following linear regression model was fitted on the data of every participant: where EEG_R2_ represents z-scored posterior response 2-locked EEG activity, EEGR1 represents z-scored posterior response 1-locked EEG activity, IRI represents the z-scored inter-response interval of the current trial. All z-scores were computed within-subject (i.e., using subject-specific across-trial mean and standard deviations) and βi represent within-subject regression coefficients. Participants’ regression coefficients were subsequently standardized by their SDs and tested against zero by means of Student’s t-tests. Due to the large number of the resulting t-tests, we corrected for multiple comparisons by means of a cluster-based permutation test with 100.000 permutations, a cluster inclusion threshold of *p* = .01 and an output threshold of *p* = .05, utilizing the Mass Univariate ERP Toolbox (Groppe, Urbach, & Kutas, 2011).

## Acknowledgements

This work was supported by a grant within the Priority Program, SPP 1772 from the German Research Foundation (Deutsche Forschungsgemeinschaft, DFG, grant number STE 1708/4-1).

## Notes

### Competing Interest Statement

The authors have declared no competing interest.

### Summary of Updates

Revision as requested by reviewers

